# Cas13b is a Type VI-B CRISPR-associated RNA-Guided RNase differentially regulated by accessory proteins Csx27 and Csx28

**DOI:** 10.1101/092577

**Authors:** Aaron Smargon, David B.T. Cox, Neena Pyzocha, Kaijie Zheng, Ian M. Slaymaker, Jonathan S. Gootenberg, Omar A. Abudayyeh, Patrick Essletzbichler, Sergey Shmakov, Kira S. Makarova, Eugene V. Koonin, Feng Zhang

## Abstract

CRISPR-Cas adaptive immune systems defend microbes against foreign nucleic acids via RNA-guided endonucleases. Using a computational sequence database mining approach, we identify two Class 2 CRISPR-Cas systems (subtype VI-B) that lack Cas1 and Cas2 and encompass a single large effector protein, Cas13b, along with one of two previously uncharacterized associated proteins, Csx27 or Csx28. We establish that these CRISPR-Cas systems can achieve RNA interference when heterologously expressed. Through a combination of biochemical and genetic experiments, we show that Cas13b processes its own CRISPR array with short and long direct repeats, cleaves target RNA, and exhibits collateral RNase activity. Using an *E. coli* essential gene screen, we demonstrate that Cas13b has a double-sided protospacer-flanking sequence and elucidate RNA secondary structure requirements for targeting. We also find that Csx27 represses, whereas Csx28 enhances, Cas13b-mediated RNA interference. Characterization of these CRISPR systems creates opportunities to develop tools to manipulate and monitor cellular transcripts.

## INTRODUCTION

CRISPR-Cas (*c*lustered *r*egularly *i*nterspaced *s*hort *p*alindromic *r*epeats and *C*RISPR-*as*sociated proteins) systems are divided into two classes, Class 1 systems, which utilize multiple Cas proteins and CRISPR RNA (crRNA) to form an effector complex, and the more compact Class 2 systems, which employ a large, single effector with crRNA to mediate interference (Makarova et al., 2015). CRISPR-Cas systems display a wide evolutionary diversity, involving distinct protein complexes and different modes of operation, including the ability to target RNA (Hale et al.,2009; Hale et al., 2012; Staals et al., 2013; Tamulaitas et al., 2014; Staals et al., 2014; Jiang et al., 2016; Abudayyeh et al., 2016; East-Seletsky et al., 2016).

Computational sequence database mining for diverse CRISPR-Cas systems has been carried out by searching microbial genomic sequences for loci harboring the *cas1* gene, the most highly conserved *cas* gene involved in the adaptation phase of CRISPR immunity (Marraffini L.A., 2015). Among other findings, this approach led to the discovery of the Class 2 subtype VI-A system with its signature effector Cas13a (previously known as C2c2), which targets RNA (Shmakov et al., 2015; Abudayyeh et al., 2016; East-Seletsky et al., 2016; Shmakov et al., *in press*). Since distinct variants of functional Class 1 CRISPR systems have been characterized that lack *cas1* (Makarova et al., 2015), we sought to identify Class 2 CRISPR-Cas systems lacking *cas1* by modifying the computational discovery pipeline so it is not seeded on Cas1. Here we report the characterization of a new Class 2 subtype, VI-B, which was discovered through this computational approach, and demonstrate that the VI-B effector, Cas13b, is an RNA-guided RNase.

## RESULTS

### Computational discovery of Class 2 subtype VI-B CRISPR systems

We designed a computational pipeline to search specifically for putative CRISPR-*cas* loci lacking Cas1 and Cas2 (Figure 1A). Fully assembled microbial genomes were searched for all proteins within 10kb of CRISPR arrays. The list of identified loci was further narrowed down using the following criteria: no more than one neighboring protein larger than 700aa (to eliminate Class 1 system false positives), presence of a putative single effector of size 900aa to 1800aa (informed by the size distribution of previously classified Class 2 effectors), and absence of *cas1* and *cas2* genes within 10kb of the CRISPR array (Experimental Procedures). Candidate effectors were grouped into families according to homology (Altschul et al., 1997; Camacho et al., 2009; Hildebrand et al., 2009), and discarded if they matched previously identified CRISPR-Cas systems (Makarova et al., 2015). To focus on likely functional CRISPR loci, we limited the candidate list to families of at least 10 non-redundant effectors in which the putative effector was near a CRISPR array for at least 50% of the members.

**Figure 1.**
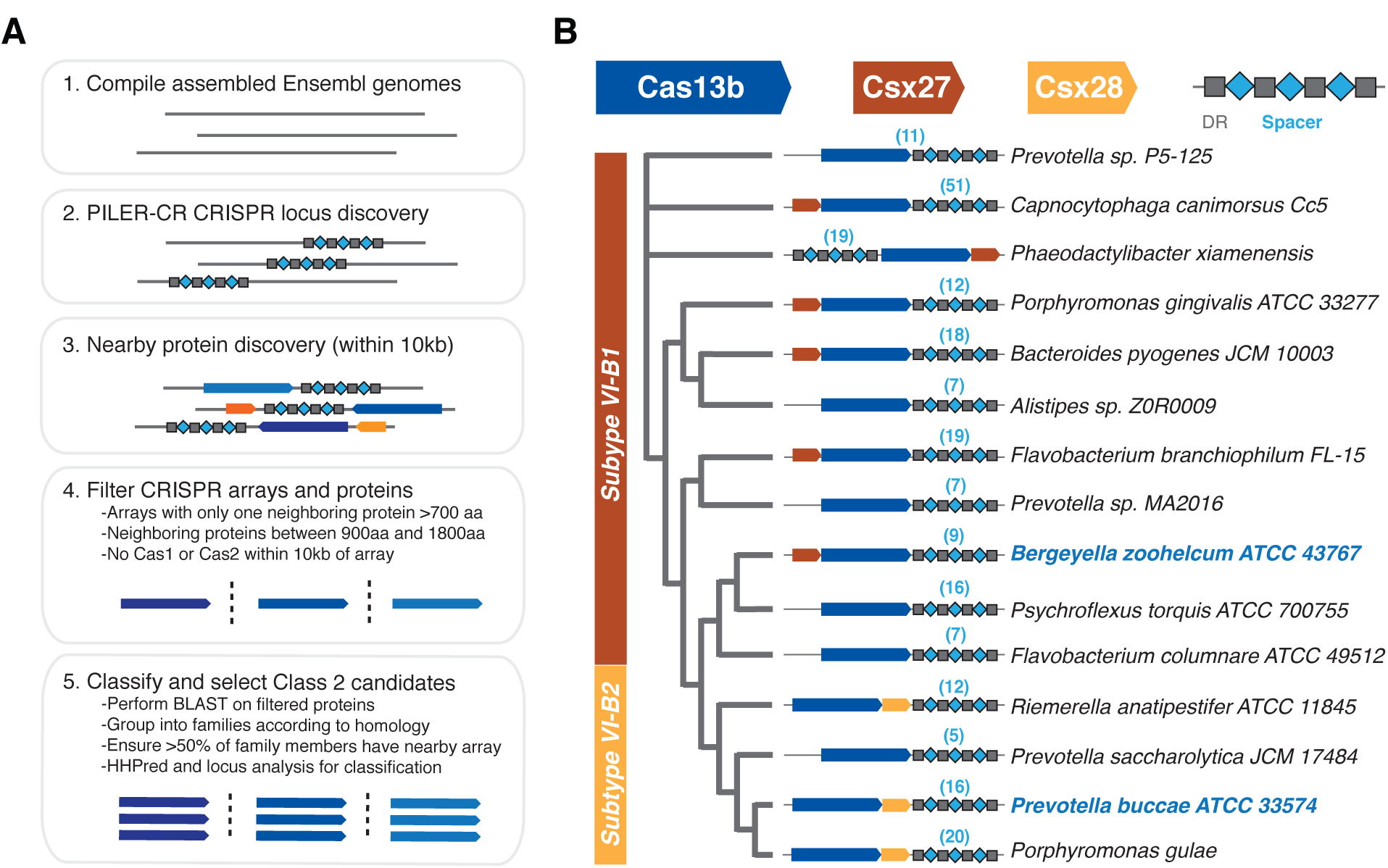
Discovery of two Class 2 CRISPR-Cas systems, subtype VI-B1 and VI-B2, containing Cas13b. **(A)** Bioinformatic pipeline to discover putative Class 2 CRISPR loci lacking Cas1 and Cas2. **(B)** A schematic phylogenetic tree of the subtype VI-B loci. Loci with *csx27* (brown) comprise variant VI-B1; loci with *csx28* (gold) comprise variant VI-B2.

Among the candidates, we identified two genetically diverse putative Class 2 CRISPR-Cas systems (105 genomic loci, 81 of which contain a unique entry Cas13b in the non-redundant NCBI protein database, and 71 of these 81 contain an annotated CRISPR array) represented in Gram-negative bacteria (Figure S1A). For several genera, in particular *Porphyromonas* and *Prevotella,* Cas13b proteins are encoded in several unique sequenced loci, and occasionally, in the same sequenced genome. These systems often co-occur with other CRISPR-Cas systems, but we identified two genomes, *Aaludibacter propionicigenes* and *Prevotella intermedia,* in which CRISPR-Cas13b was the only detectable CRISPR locus (Figure S1).

All these loci encode a large (~1100aa) candidate effector protein and, in about 80% of the cases, an additional small (~200aa) protein (Figure 1B, S1). The putative effector proteins contain two predicted HEPN domains (Anantharaman et al., 2013) at their N- and C-termini (Figure S2A), similar to the domain architecture of the large effector of subtype VI-A (Cas13a) (Shmakov et al., 2015). Beyond the occurrence of two HEPN domains, however, there is no significant sequence similarity between the predicted effector and Cas13a. These systems were also identified by a generalized version of the pipeline described above as part of a comprehensive analysis of Class 2 CRISPR-Cas systems, and were classified into subtype VI-B, with predicted effector protein Cas13b (Shmakov et al., in press).

### CRISPR-Cas13b loci contain small accessory proteins

The identity of the putative accessory protein correlates with the two distinct branches in the phylogenetic tree of Cas13b (Figure 1B, S1), indicative of the existence of two variant systems, which we denote VI-B1 (accessory protein referred to as Csx27) and VI-B2 (accessory protein referred to as Csx28). While subtype VI-B2 systems almost invariably contain *csx28, csx27* is less consistently present in VI-B1 loci. The protein sequences of Csx27 and Csx28 show no significant similarity to any previously identified Cas proteins. Both Csx27 and Csx28 were predicted to contain one or more transmembrane segments (Figure S3A). However, Csx27 of *Bergeyella zoohelcum* and Csx28 of *Prevotella buccae* tagged with RFP at either the N- or C-terminus did not show membrane localization when expressed in *E. coli* (Figure S3B). In addition to the predicted hydrophobic domains, analysis of the multiple sequence alignment of Csx28 proteins indicated the presence of a divergent HEPN domain (Figure S2).

### Cas13b-associated CRISPR arrays display unique features

In contrast to the distinct putative accessory proteins, both variants of subtype VI-B systems show distinct, conserved features in the CRISPR arrays. The direct repeats in the CRISPR arrays are conserved in size, sequence, and structure, with a length of 36 nt, a poly-U stretch in the open loop region, and complementary sequences 5’-GUUG and CAAC-3’ at the ends of the repeat predicted to yield a defined secondary structure mediated by intramolecular base-pairing (Figure S4, S5A). Our analysis revealed 36 Cas13b spacers mapped with greater than 80% homology to unique protospacers in phage genomes. Twenty-seven of the identified Cas13b spacers targeted the coding strand of phage mRNA, while 7 spacers targeted the noncoding strand and 2 spacers targeted regions of the phage genome without predicted transcripts. Although the composite of these imperfect mappings revealed no consensus flanking region sequence (Figure S5B) (Biswas et al., 2013), the well-conserved protospacer length of 30 nt, combined with the conserved direct repeat sequence and length, suggests that the nucleic acid targeting rules may be similar among different VI-B loci.

RNA sequencing of the total RNA from *B. zoohelcum* (subtype VI-B1) showed processing of the pre-crRNA into a 66-nt mature crRNA, with the full 30-nt 5’ spacer followed by the full 36-nt 3’ direct repeat (Figure 2A). A longer 118-nt crRNA, distal to the 36-nt crRNAs in the CRISPR array and consisting of fragments of 5’ and 3’ direct repeat sequences interrupted by an intervening repeat sequence, was also processed. This phenomenon was computationally predicted to occur in other VI-B loci, such as those from *Capnocytophaga canimorsus*, *Myroides odoratimimus*, and *Riemerella anatipestifer*. Other CRISPR Class 2 effectors are known to process their arrays without involvement of additional RNases (Zetsche et al., 2015; East-Seletsky et al., 2016). Similarly, we find that purified BzCas13b is capable of cleaving its CRISPR array, generating mature crRNAs with short or long direct repeats, and spacers which are not further processed beyond 30nt, an activity which is not affected by mutation of the predicted catalytic residues of the HEPN domain (Figure 2B, S6).

**Figure 2.**
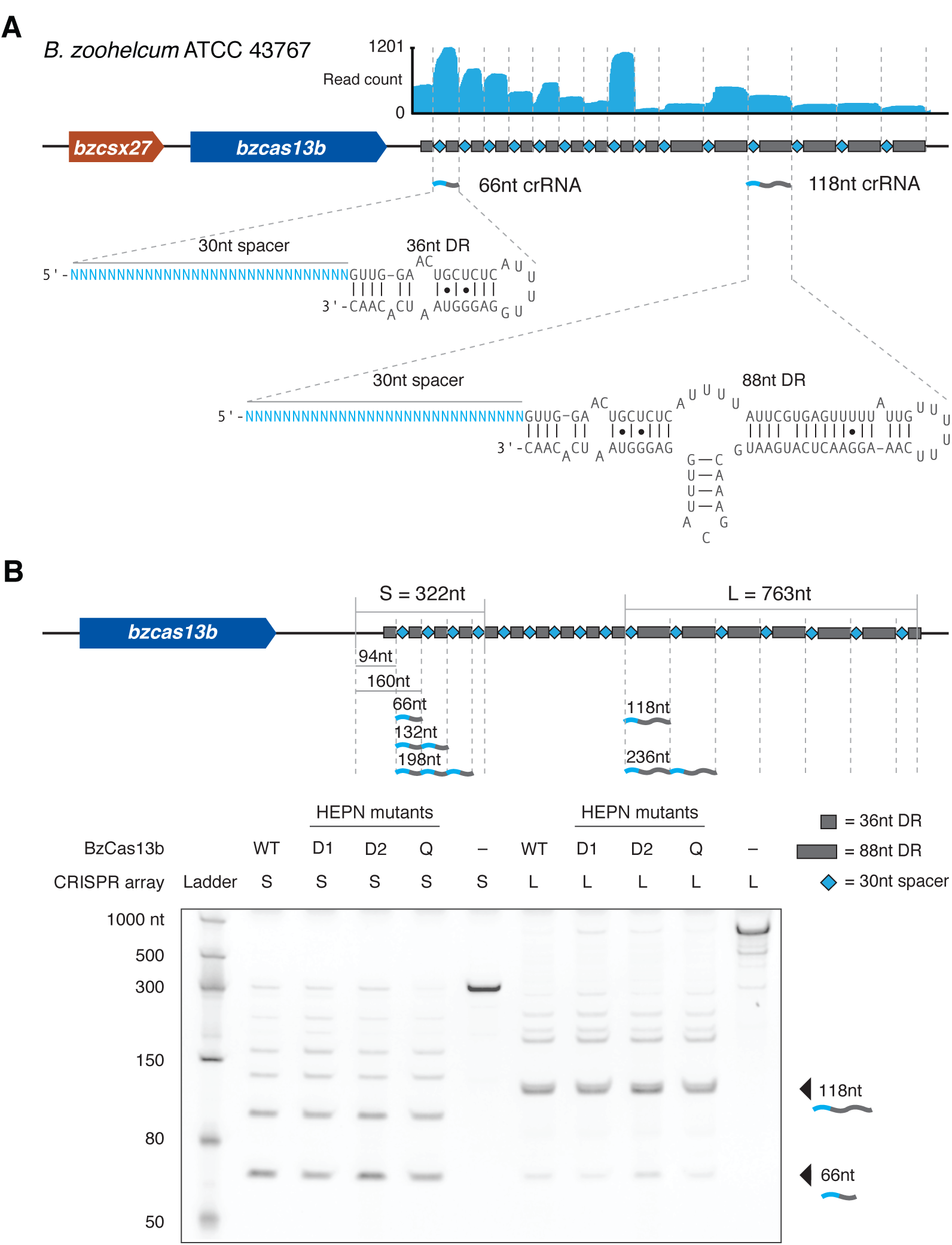
Cas13b from the VI-B1 locus processes a CRISPR array with two direct repeat variants. **(A)** RNA-Sequencing of the native VI-B1 locus from *Bergeyella zoohelcum* ATCC 43767. **(B)** Denaturing gel showing cleavage products of in vitro synthesized short-DR-or long-DR-containing CRISPR arrays from the *B. zoohelcum* genome by either wildtype or HEPN mutant BzCas13b (D1, R116A/H121A; D2, R1177A/H1182A; Q, R116A/H121A/R1177A/ H1182A). The schematic shows fragment lengths of a cleaved CRISPR array.

### An *E. coli* essential gene screen reveals targeting rules for *BzCas13b*

To validate the expected interference activity of the VI-B system and to determine the targeting rules for the VI-B1 locus from *B. zoohelcum,* we developed an *E. coli* essential gene screen (Figure 3A). For this negative selection screen, we generated a library of 54,600 unique spacers tiled with single-nucleotide resolution over the coding region of 45 monocistronic essential genes (Gerdes et al., 2003; Baba et al., 2006), plus 60 nt into the 5’ and 3’ UTRs. We also included 1100 randomly generated non-targeting spacers to establish baseline activity. We then transformed this library with plasmids carrying *bzcas13b (cas13b* gene from *B. zoohelcum)* and *bzcsx27,* just *bzcas13b,* or a control empty vector. After quality-control-filtering of all screened spacers (Experimental Procedures), we found a statistically significant depletion of targeting spacers over non-targeting spacers, indicating that Cas13b, alone or with Csx27, can achieve nucleic acid interference (Figure 3B).

**Figure 3.**
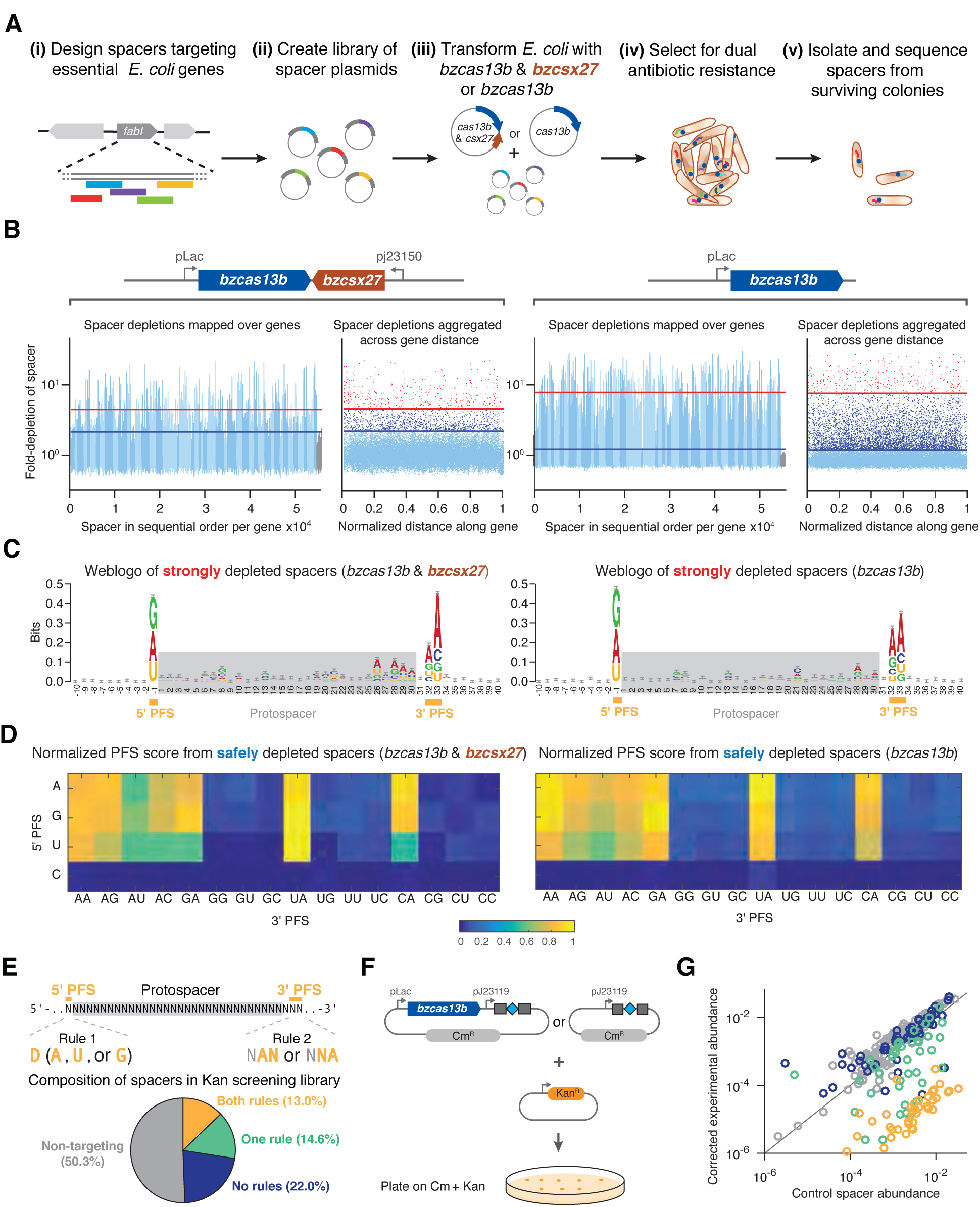
Heterologous expression of Cas13b mediates knockdown of *E. coli* essential genes by a double-sided PFS. **(A)** Design of *E. coli* essential gene screen to determine targeting rules of nucleic acid interference. **(B)** Manhattan plots of mean spacer depletions mapped over 45 genes and aggregated across normalized gene distance for either the full *B. zoohelcum* VI-B1 locus (left) or *cas13b* alone (right), with non-targeting spacers in gray, safely depleted spacers (>5σ above mean depletion of non-targeting spacers) above blue line, and strongly depleted spacers (top 1% depleted) above red line. For the full locus, 36,142 targeting spacers and 630 non-targeting spacers passed QC filter. Of the targeting, 367 are strongly depleted and 1672 are safely depleted. For *cas13b* alone, 35,272 targeting spacers and 633 non-targeting spacers passed QC filter. Of the targeting, 359 are strongly depleted and 6374 are safely depleted. **(C)** Weblogo of sequence motifs of strongly depleted *B. zoohelcum* spacers. **(D)** Normalized PFS score matrix, where each score is the ratio of number of safely depleted *B. zoohelcum* spacers to total number of spacers for a given PFS, scaled so that maximum PFS score is 1. **(E)** Spacers targeting kanamycin to validate PFS targeting rules of 5’ PFS (D) and 3’ PFS (NAN or NNA). **(F)** Schematic of kanamycin validation screen for *B. zoohelcum cas13b* in *E. coli.* **(G)** Results from kanamycin validation screen; spacer abundances versus control for individual *B. zoohelcum* spacers, with abundances colored by type of spacer.

To assess the targeting rules for Cas13b, we established two spacer depletion levels: strongly depleted (top 1% of depleted spacers) and safely depleted (spacers depleted 5σ above the mean depletion of the filtered non-targeting spacers). From spacers passing the strongly depleted cutoff we derived sequence motifs qualitatively identifying a double-sided protospacer flanking sequence (PFS) (Figure 3C). Because each position in a sequence weblogo is assumed to be independent, we developed a more quantitative, base-dependent PFS score defined as the ratio of the number of safely depleted spacers to the number of all spacers with a given PFS, normalized across all PFS scores (Figure 3D).

The normalized PFS scores revealed a 5’ PFS of D (A, U, or G) and 3’ PFS of NAN or NNA, consistent for Cas13b with Csx27, as well as for Cas13b alone. To validate these sequence-targeting rules, we performed an orthogonal depletion screen with Cas13b alone, targeting the Kanamycin resistance gene (Figures 3E, 3F). Four classes of spacers were created: non-targeting, targeting with both 5’ and 3’ PFS rules, targeting with only the 5’ or 3’ PFS rule, and targeting with neither rule. Consistent with our findings from the *E. coli* essential gene screen, the 5’ and 3’ PFS spacers resulted in the highest Kanamycin sensitivity (Figures 3G, S7).

### BzCas13b cleaves single-stranded RNA and exhibits collateral activity in vitro

Based on the presence of the computationally predicted HEPN domains that function as RNases in other CRISPR-Cas systems, including VI-A and some Class 1 systems (Kim et al., 2013; Staals et al., 2014; Sheppard et al., 2016; Abudayyeh et al., 2016), we anticipated that Cas13b interferes with RNA. We confirmed this by demonstrating that purified Cas13b exclusively cleaves single-stranded RNA with both direct repeat architectures (Figure 4A, S8A). We then validated the PFS targeting rules biochemically, showing that a 5’ PFS of C greatly inhibits single-stranded RNA cleavage (Figure 4B), whereas a 3’ PFS of NAN or NNA enhances this activity (Figure 4C).

**Figure 4.**
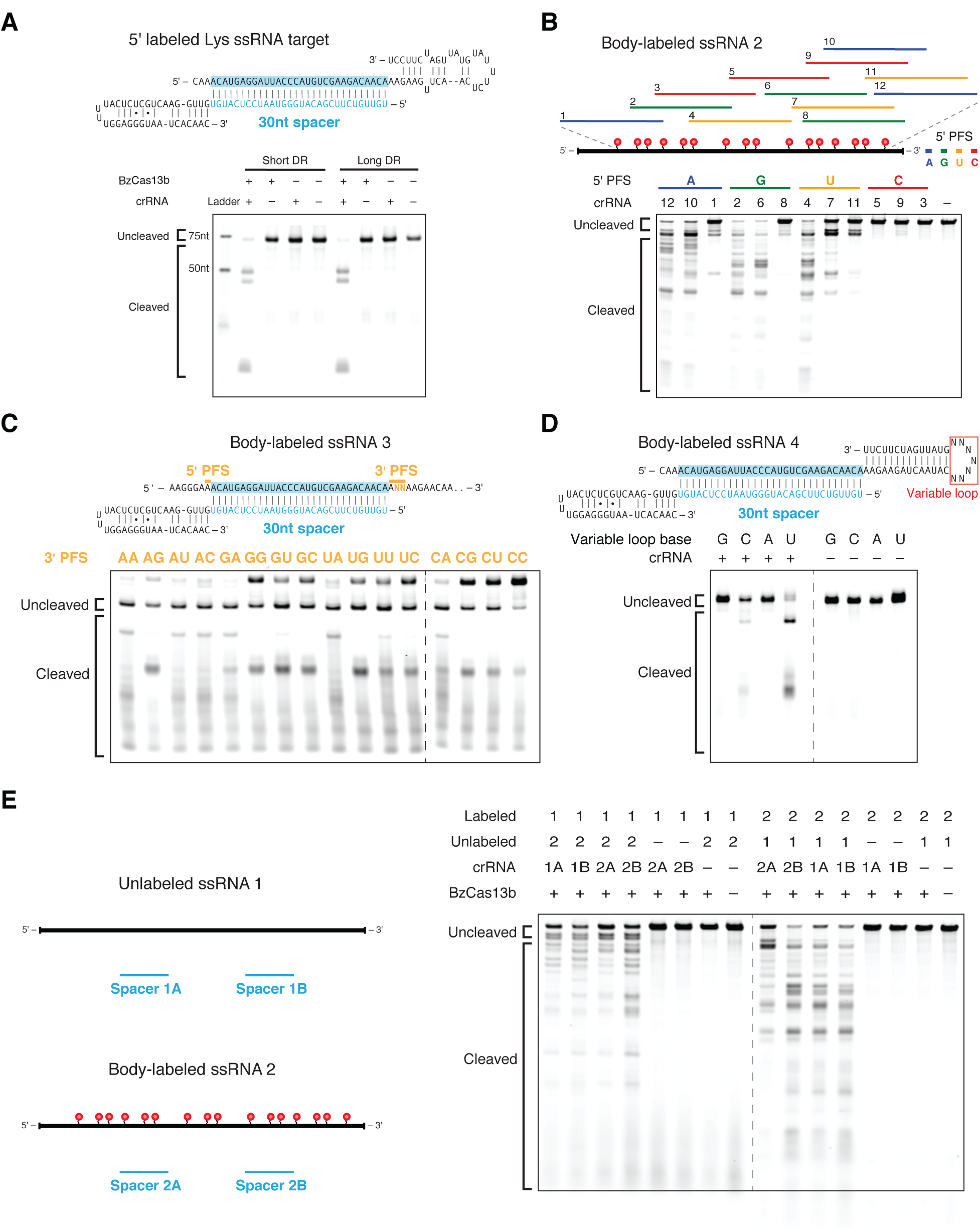
Cas13b is a programmable single-stranded RNase with collateral activity. **(A)** Schematics showing the RNA secondary structure of the cleavage target alone (left top), cleavage target in complex with a targeting 30nt spacer (left middle), and crRNAs with either the short or long direct repeat (left bottom). For the cleavage target alone, the dark bases show the spacer-targeted region. Numbers along the targets represent nucleotide distance from the 5’ labeled end. Denaturing gel demonstrating short direct repeat and long direct repeat crRNA-mediated ssRNA cleavage (right). Reactions were incubated for 10 minutes. The short and long direct repeat crRNAs cleave ssRNA with similar efficiency. The ssRNA target is 5’ labeled with IRDye 800. Three cleavage sites are observed. **(B)** Schematic showing three numbered protospacers for each colored 5’ PFS on a body-labeled ssRNA target (top). Denaturing gel showing crRNA-guided ssRNA cleavage activity demonstrating the requirement for a D 5’ PFS (not C) (bottom). Reactions were incubated for 60 minutes. crRNAs correspond to protospacer numbered from the 5’ to the 3’ end of the target. **(C)** Schematic of a body-labeled ssRNA substrate being targeted by a crRNA (top). The protospacer region is highlighted in blue, and the 5’ PFS and 3’ PFS sequences are indicated by the gold bars. The gold letters represent the altered sequences in the experiment. Denaturing gel showing crRNA-guided ssRNA cleavage activity demonstrating the preference for the NAN or NNA 3’ PFS after 60 minutes of incubation (bottom). The 5’ PFS is tested as A, and the 3’ PFS is tested as ANN. The gold 3’ PFS letters represent the RNA bases at the second and third 3’ PFS position within each target ssRNA. **(D)** Schematic showing the secondary structure of the body labeled ssRNA targets used in the denaturing gel. The variable loop of the schematic (represented as N^7^) is substituted with seven monomers of the variable loop base in the gel (top). Denaturing gel showing cleavage bands by variable loop base (bottom). The targets were incubated for 30 minutes. **(E)** Denaturing gel showing BzCas13b collateral cleavage activity after 30 minutes of incubation, with schematic of cleavage experiment to the right. Two crRNAs (A and B) target substrate 1 (1A and 1B) or substrate 2 (2A and 2B).

Other HEPN domain-containing CRISPR-Cas RNA-targeting systems, such as Csx1 from the Type III-B CRISPR-Cas systems, preferentially cleave targets containing specific singlestranded nucleotides (Sheppard et al., 2016). To determine if Cas13b exhibits such a preference, we tested an RNA substrate with a variable homopolymer loop outside of the spacer:protospacer duplex region (Figure 4D). We observed cleavage at pyrimidine residues, with a strong preference for uracil. This activity is abolished in the presence of EDTA (Figure S8B), suggesting a divalent metal ion-dependent mechanism for RNA cleavage akin to that of a similar HEPN-containing, Class 2 effector protein, Cas13a (Abudayyeh et al, 2016; East-Seletsky et al., 2016).

Given that Cas13a has also been reported to cleave RNA non-specifically once activated by interaction with the target (”collateral effect”) (Abudayyeh et al., 2016; East-Seletsky et al.,2016) we sought to test the ability of Cas13b to cleave a second, non-specific substrate following target cleavage. Using an in vitro assay similar to the one we previously used with Cas13a (Abudayyeh et al., 2016), we incubated Cas13b-crRNA complexes with both a target and non-target RNA substrate. We observed collateral cleavage of the non-targeted RNA, but only in the presence of the target RNA (Figure 4E).

### Cas13b shows robust HEPN-dependent interference and is repressed by Csx27 activity

To validate RNA interference in vivo, we assayed interference against the lytic, single-stranded RNA bacteriophage MS2, whose life cycle contains no DNA intermediates. We performed an MS2 drop plaque assay at serial dilutions of phage for both *bzcas13b* with *bzcsx27* and *bzcas13b* alone with three spacers targeting the MS2 genome, two at the *lys-rep* interface and one in *rep,* as well as one non-targeting spacer (Figure 5A). We observed substantial reduction in plaque formation for all targeting spacers compared to the non-targeting spacer, confirming sequence-specific RNA targeting by VI-B1 systems. (Figure 5A, S9). Notably, the presence of *bzcsx27* weakened RNA interference by *bzcas13b* for all three spacers.

To confirm the lack of DNA interference in vivo, we modified an existing plasmid interference assay with a protospacer placed either in-frame at the 5’ end of the *bla* ampicillin-resistance gene (transcribed target) or upstream of the *bla* gene promoter (non-transcribed target). Bacteria cotransformed with *bzcas13b* and spacer as well as the non-transcribed target plasmid survived at a comparable rate to co-transformation of the same target with the empty vector on dual antibiotic selection. For bacteria co-transformed with the transcribed target, the colony forming unit rate under dual antibiotic selection was reduced by ~2 orders of magnitude in the presence of *bzcas13b,* corroborating that Cas13b exclusively targets RNA in vivo (Figure 5B).

**Figure 5.**
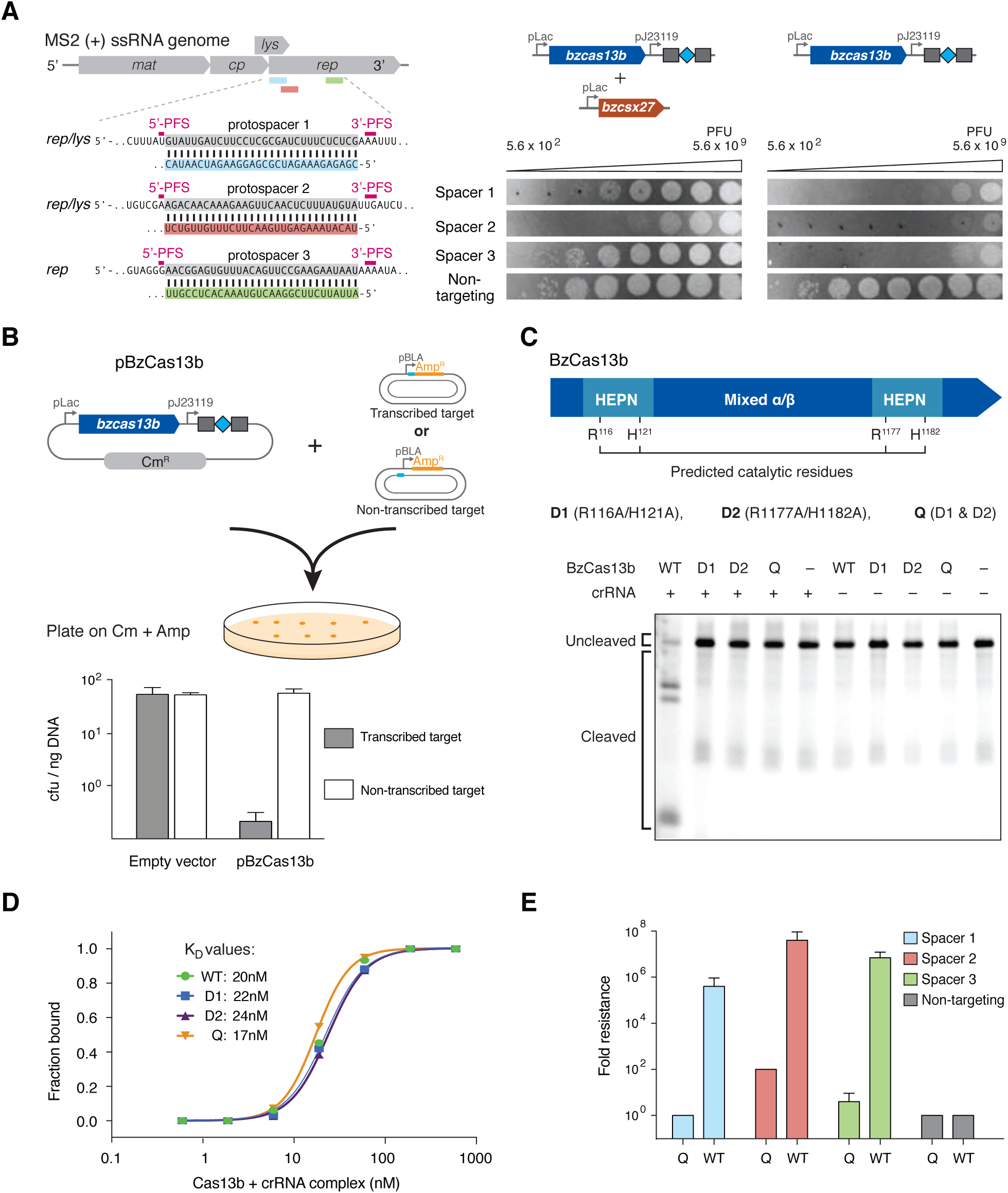
HEPN domains mediate RNA cleavage by Cas13b, whose activity is repressed by Csx27. **(A)** Protospacer design for MS2 phage plaque drop assay to test RNA interference (left). Plaque drop assay for full *B. zoohelcum* VI-B1 locus (center) and *bzcas13b* (right). **(B)** DNA interference assay schematic and results. A target sequence is placed in frame at the start of the transcribed *bla* gene that confers ampicillin resistance or in a non-transcribed region of the same target plasmid. Target plasmids were co-transformed with *bzcas13b* plasmid or empty vectors conferring chloramphenicol resistance and plated on double selection antibiotic plates. **(C)** Denaturing gel showing ssRNA cleavage activity of WT and HEPN mutant BzCas13b. The protein and targeting crRNA complexes were incubated for 10 minutes. **(D)** Electrophoretic Mobility Shift Assay (EMSA) graph showing the affinity of BzCas13b proteins and targeting crRNA complex to a 5’ end labeled ssRNA. The EMSA assay was performed with supplemental EDTA to reduce any cleavage activity. **(E)** Quantification of MS2 phage plaque drop assay with *B. zoohelcum* wildtype and R116A/H121A/R1177A/H1182A Cas13b.

We next tested if predicted catalytic residues in the HEPN domains were responsible for RNA cleavage by Cas13b. Three HEPN mutants were obtained by replacing the conserved catalytic arginines and histidines in the two HEPN domains with alanines (R116A/H121A, termed domain 1 (D1); R1177A/H1182A, termed domain 2 (D2); and R116A/H121A/R1177A/H1182A, termed quadruple (Q)) (Figure S6). All mutants lacked observable cleavage activity (Figure 5C), yet retained RNA binding capacity in vitro (Figure 5D, S10A). The wildtype and all three HEPN mutant Cas13b proteins showed comparable binding affinities for a single-stranded target RNA substrate, with K_D_ values ranging from 17nM to 24nM (Figure 5D, S10A). The K_D_ for off-target binding was found to be greater than 188nM (Figure S10B).

We confirmed the involvement of the HEPN domains in RNA interference in vivo, finding ~5.5 orders of magnitude decrease in resistance to MS2 phage in the quadruple HEPN mutants versus wildtype Cas13b (Figures 5E, S9). Interestingly, quadruple mutant Cas13b with spacers 2 and 3 still showed weak phage resistance, potentially due to catalytically inactive Cas13b binding to phage genomic RNA, leading to reduced phage replication.

### Computational modeling predicts additional targeting rules governing *Cas13b*

Our sequence-based targeting results from the *E. coli* essential gene screen implied the existence of additional RNA-targeting rules beyond the PFS (only ~18% of spacers were safely depleted for *bzcas13b;* from the PFS rules alone, the expected value would be ~33%). Given that RNA targets contain a variety of secondary structures, we sought to determine how RNA accessibility impacts targeting. Using the Vienna RNAplfold method (Bernhart et al., 2006), which has been successfully employed to predict RNAi efficiency (Tafer et al., 2008) (Figure 6A), we trained and tested an RNA accessibility model for spacer efficiency on our screen data, and found that RNA accessibility matters the most in the protospacer region most distal to the direct repeat of the crRNA (Figures 6B, 6C).

**Figure 6.**
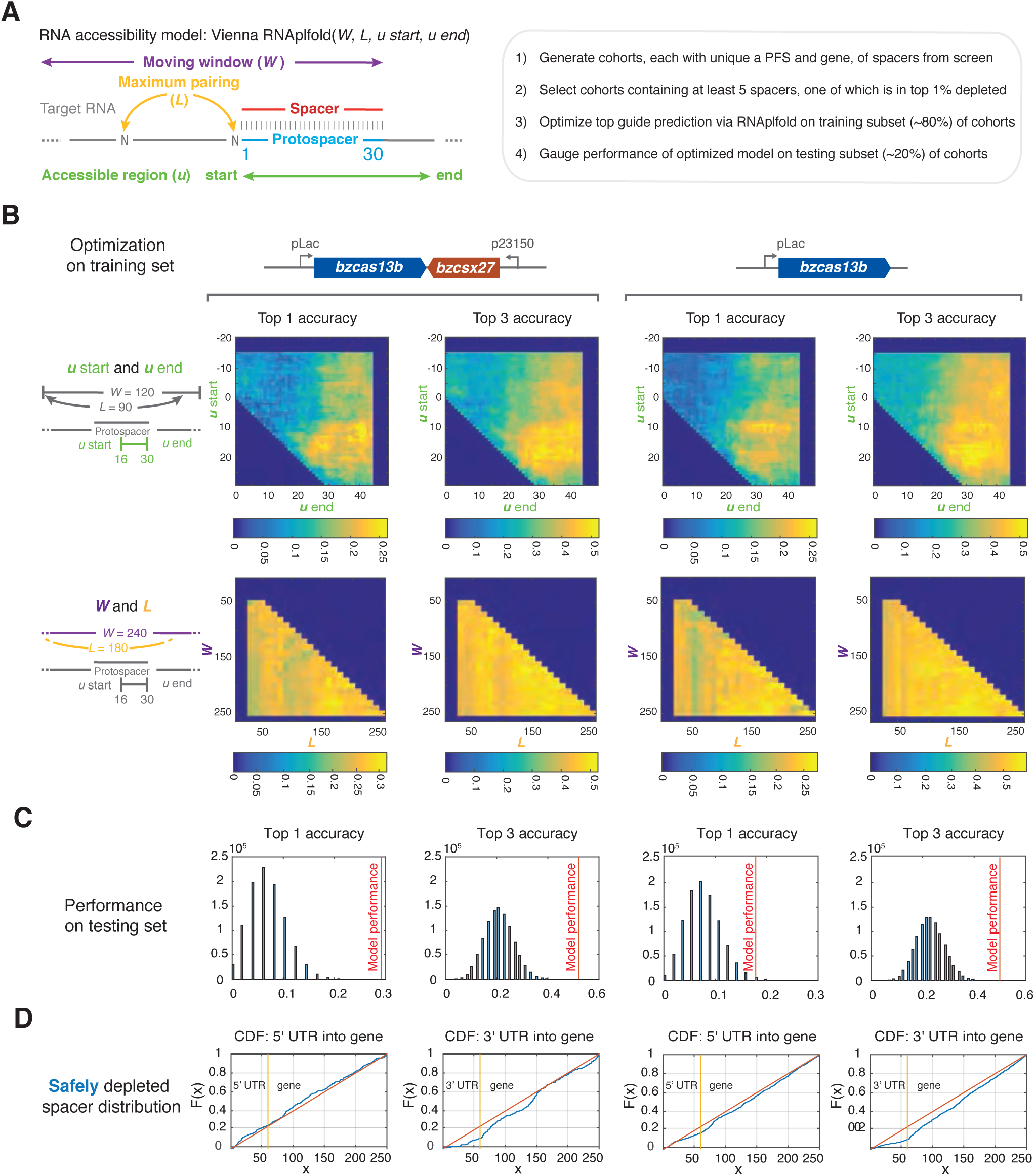
Efficient RNA targeting by Cas13b is correlated with local RNA accessibility. **(A)** Methodology of secondary structure-mediated spacer efficiency analysis of *E. coli* essential gene screen data with Vienna RNAplfold. **(B)** Optimization of *top 1 accuracy* (computationally predicted most accessible spacer matches the top experimentally depleted spacer) and *top 3 accuracy* (computationally predicted top spacer falls in top 3 experimentally depleted spacers) on randomly selected *B. zoohelcum* training dataset using RNAplfold, first with *u start* and *u end*, and then with *W* and *L*. **(C)** Performance of optimized RNAplfold model on randomly selected *B. zoohelcum* testing dataset (48 cohorts for full *B. zoohelcum* VI-B1 locus, 56 cohorts for *bzcas13b)* against 10^6^ Monte Carlo simulations: empirical P-values from left to right of 3e-6, 1e- 6, 8.7e-3, 6e-6. **(D)** Empirical cumulative distribution of safely depleted *B. zoohelcum* spacers over all genes from 5’ UTR into gene and from 3’ UTR into gene. Yellow line separates UTR and gene, red line is theoretical cumulative distribution function of uniformly distributed spacers, and blue line is empirical cumulative distribution of safely depleted *B. zoohelcum* spacers.

Given the collateral activity observed in vitro, we examined our screen data for indications of non-specific RNA cleavage by Cas13b. To this end, we calculated the empirical cumulative distribution functions of safely depleted spacers aggregated across all essential genes from the 5’ UTR into the gene and from the 3’ UTR into the gene (Figure 6D). Because cleavage closer to the 5’ UTR is more likely to disrupt gene function, without non-specific RNase activity, we would expect an overrepresentation of spacers in the 5’ UTR and an underrepresentation in the 3’ UTR. By contrast, in the presence of collateral activity a nearly uniform distribution would be expected. From our screen data, we observed only a marginal underrepresentation of spacers in the 3’ UTR compared to a uniform distribution, suggesting that collateral activity may occur in vivo.

### CRISPR-Cas13b effectors are differentially regulated by Csx27 and Csx28

To determine if the established RNA targeting rules generalize across the subtype VI-B systems from diverse bacteria, we characterized the subtype VI-B2 locus from *P. buccae.* RNA sequencing of the CRISPR array revealed processing effectively identical to that of *B. zoohelcum,* excluding the long crRNA (Figure S11). The *E. coli* essential gene screen with *pbcas13b* and *pbcsx28* or *pbcas13* alone led to the identification of a PFS matrix similar to that of *B. zoohelcum,* with certain PFS’s disfavored (Figures 7A, S12). Similar to BzCas13b, PbCas13b was found to cleave targeted single-stranded RNA in vitro (Figure S8C). As with *bzcsx27,* the presence of *pbcsx28* did not appreciably alter the PFS. We also repeated the secondary structure analysis with *pbcas13b,* and a comparable RNAplfold model applied (Figures S13A, S13B). Strikingly, in these experiments, the safely depleted spacers for *pbcas13b* alone were highly biased to the beginning of the 5’ UTR of genes, suggestive of inhibited or more spatially localized RNase activity in the absence of *pbcsx28* (Figure S13C). We further explored the apparent reduced activity of *pbcas13b* alone relative to the respective full CRISPR-Cas locus using the MS2 phage plaque drop assay and found that *pbcsx28* enhances MS2 phage interference by up to four orders of magnitude (Figures 7B, S9). The differential ability of *csx27* to repress and *csx28* to enhance *cas13b* activity generalizes across thousands of spacers in the *E. coli* essential gene screen (Figure 7C) highlighting the distinctive regulatory modes of the two variants of subtype VI-B CRISPR-Cas systems.

**Figure 7.**
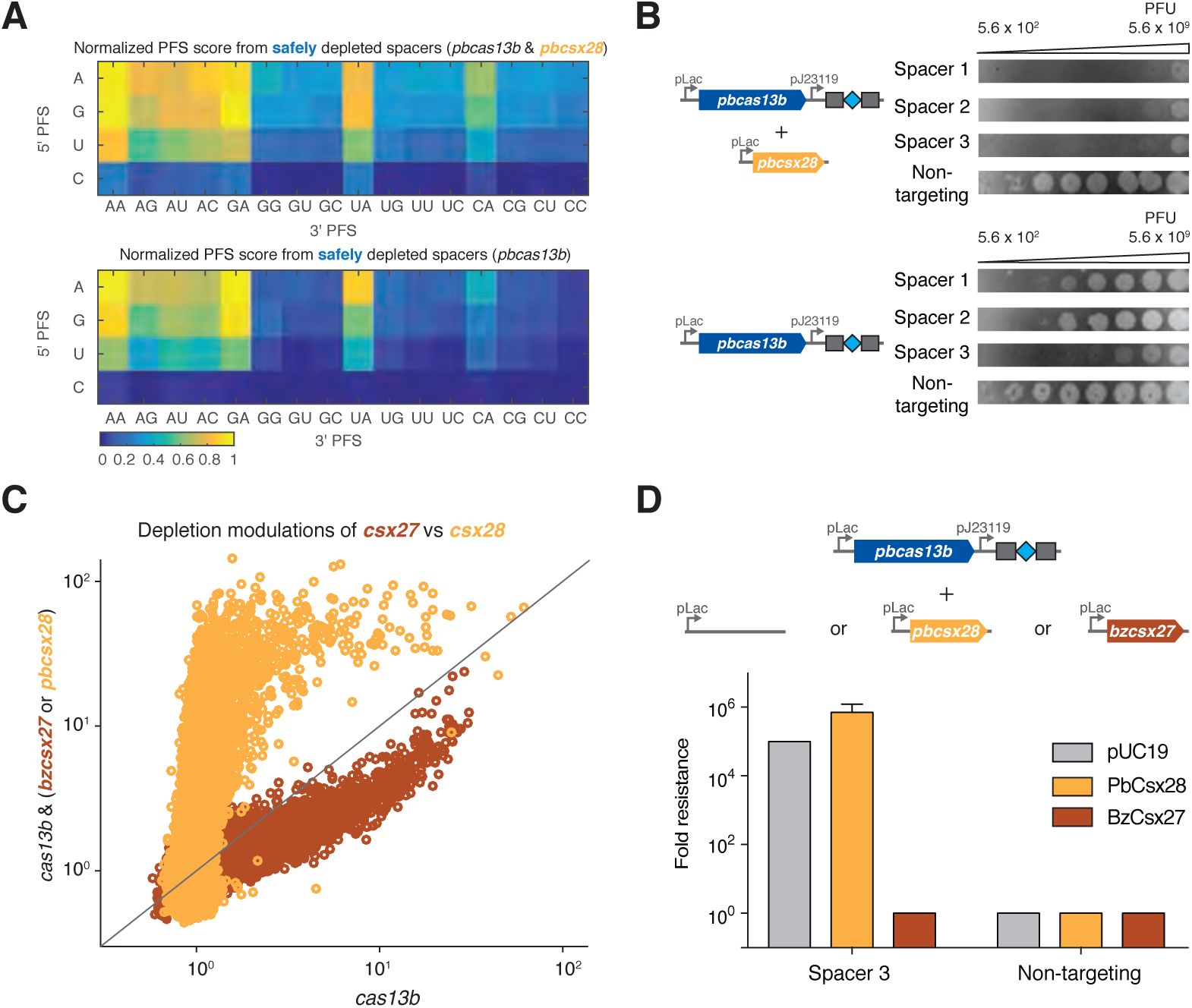
Class 2 type VI-B systems are differentially regulated across two loci by Csx27 and Csx28. **(A)** Normalized PFS matrix, for *P. buccae* VI-B2 locus (left) and *pbcas13b* (right). **(B)** MS2 Plaque drop assay for full *P. buccae* VI-B2 locus (left) and *pbcas13b* (right). **(C)** Spacer depletions of *bzcas13b* with and without *bzcsx27* (brown), as compared to *pbcas13b* with and without *pbcsx28* (gold). **(D)** Plaque assay images for cells co-transformed with *pbcas13b* and the indicated *csx* expression plasmid. In the rightmost column are cells co-transformed with pUC19 empty vector.

To further explore the ability of the small accessory proteins to modulate Cas13b activity, we tested if Csx27 can also repress PbCas13b using the MS2 drop plaque assay. Cells cotransformed with *pbcas13b* and *bzcsx27* expression plasmids exhibited a 10^5^ fold reduction in interference activity relative to pUC19 empty vector, suggesting that Csx27 exerts an inhibitory effect on PbCas13b (Figures 7D, S9). The ability of Csx27 to modulate the interference activity of BzCas13b and PbCas13b suggests that it is a modular protein that can function across multiple VI-B loci (Figure 7E).

## DISCUSSION

Here we describe two RNA-targeting CRISPR Class 2 systems of subtype VI-B (VI-B1 and VI-B2), containing the computationally discovered RNA-guided RNase Cas13b. Type VI-B systems show several notable similarities to the recently characterized VI-A system. The single protein effectors of both systems cleave single-stranded RNA via HEPN domains, process their CRISPR arrays independent of the HEPN domains, and exhibit collateral RNase activity. Cas13b proteins, however, show only limited sequence similarity to Cas13a, and the common ancestry of the two type VI subtypes remains uncertain. Furthermore, the type VI-B systems differ from VI-A in several other ways, including the absence of both *cas1* and *cas2,* which are involved in spacer acquisition in other CRISPR-Cas systems (Mohanraju et al., 2016). The VI-B CRISPR arrays contain multiple spacers that differ among closely related bacterial strains, suggesting that acquisition does occur, either autonomously or possibly *in trans,* by recruiting Cas1 and Cas2 encoded in other CRISPR-Cas loci from the same genome. Additionally, VI-B systems differ from VI-A systems by the presence of the small accessory proteins Csx27 (VI-B1 systems) and Csx28 (VI-B2 systems), which exhibit opposing regulatory effects on Cas13b activity.

Repression of Cas13b by Csx27 in VI-B1 systems could be part of an important regulatory mechanism of phage interference. The ability of Csx27 to repress Cas13b activity may be a general property, as we found that it can also repress PbCas13b (subtype VI-B2). In the case of type VI-B2 systems, Csx28 might enhance the collateral activity of Cas13b to inactivate numerous transcripts of invading bacteriophages or to promote programmed cell death. Both Csx27 and Csx28 contain predicted long, hydrophobic α-helices that might enable them to physically interact with Cas13b, but this remains to be determined. We did not find any instances of Csx27 or Csx28 encoded outside type VI-B loci, suggesting that these proteins might function in tight cooperation with Cas13b.

As with previously characterized Class 2 CRISPR-Cas effectors, such as Cas9 and Cpf1, there is enormous potential to harness Cas13b for use as a molecular tool. A holistic understanding of the factors that affect target selection is essential to the success of any such tools, particularly those that target RNA, where secondary structure will likely impact activity. We therefore developed an *E. coli* essential gene screen to fully explore the targeting rules of Cas13b. This *E. coli* screen offers several advantages by increasing the number of guides testable in a single experiment to explore how diverse spacer and flanking sequences may affect Cas13b activity. This screen revealed a double-sided PFS in VI-B systems, which may give insight into Cas13b protein-RNA interactions, and could help improve specificity by expanding sequence targeting constraints (Ran et al., 2015).

The characterization of Cas13b and other RNA-targeting CRISPR systems raises the prospect of a suite of precise and robust in vivo RNA manipulation tools for studying a wide range of biological processes (Filipovska et al., 2011; Mackay et al., 2011; Abil et al., 2015). The ability of Cas13b to process its own CRISPR array could be extended to multiplex transcriptome engineering. In addition, the VI-B functional long direct repeats could be altered to incorporate stem loops akin to the Cas9-SAM system (Konermann et al., 2015). Like Cas9 and Cpf1, Cas13a and Cas13b may be utilized for complementary applications in science and technology.

## EXPERIMENTAL PROCEDURES

### Computational Sequence Analysis

From complete compiled Ensembl Release 27 genomes (Yates et al., 2016), CRISPR repeats were identified using PILER-CR (Edgar, 2007). Proteins within 10kb of identified CRISPR arrays were clustered into loci, with loci rejected if more than one protein of size 700 amino acids or larger or if either Cas1 or Cas2 were present. For candidate Class 2 effectors, only proteins in these remaining loci of size 900aa to 1800aa were selected. These candidate effectors were subjected to the BLASTP (Camacho et al., 2008) search against the NCBI non-redundant (NR) protein sequence database with an E-value cutoff of 1e-7. All discovered proteins were then grouped into putative families via a nearest-neighbor grouping with the same E-value cutoff. Only putative families with at least ten candidate effectors and more than 50% of candidate effectors within 10kb of CRISPR arrays were considered. HHpred (Remmert et al., 2012) and existing CRISPR locus classification rules (Makarova et al., 2015; Shmakov et al., in press) were used to classify each family, leaving Cas13b as the only unclassified family. Within this family, truncated or suspected partially sequenced effectors were discarded, leaving 105 loci, and 81 with a non-redundant protein. Multiple sequence alignments on these 81 proteins (as well as the accessory Csx27 and Csx28 proteins) were performed using BLOSUM62 (Henikoff and Henikoff, 1992) to identify the HEPN domains and to sort the loci into phylogenetic trees. Vienna RNAfold (Anantharaman et al., 2013; Lorenz et al., 2011) was used to predict the secondary structure of each direct repeat, whose transcriptional orientation was chosen as identical to that of Cas13b in its locus. CRISPRTarget (Biswas et al., 2013) was used to search the spacers in each locus against NCBI phage and plasmid genomes. Weblogos were generated for all unique direct repeats and protospacer flanking sequences (Crooks et al., 2004). TMHMM Server v. 2.0 (Moller et al., 2001) was used to predict the transmembrane helices in Csx27 and Csx28.

### RFP-Tagged Protein Fluorescent Imaging

One Shot Stbl3 Chemically Competent *E. coli* were transformed with plasmids containing RFP (negative control) or RFP fused to the N- or C-terminus of Csx27 of *B. zoohelcum* or Csx28 of *P. buccae* (Table S1). Clones were cultured up in 5mL of antibiotic LB overnight, then spun down at 5000g and resuspended in PBS with 1% methanol-free formaldehyde. After 30 minutes fixation, cells were washed once with PBS and then diluted 1:2 in PBS. 5uL of sample was pipetted onto a silane-coated slide, which was covered with a coverslip. Fluorescent imaging was performed in a 63x objective microscope with oil immersion.

### Bacterial RNA-Sequencing

RNA was isolated and prepared for sequencing using a previously described protocol (Shmakov et al., 2015; Heidrich et al., 2015). For RNA sequencing native *B. zoohelcum* ATCC 43767 grown under the same conditions, we repeated the experiment with a modified protocol, omitting TAP prior to 5’ adapter ligation, to promote enrichment of processed transcripts originating from the CRISPR array. For heterologous *P. buccae* ATCC 33574 RNA sequencing in *E. coli,* we cloned the locus into pACYC184 (Table S1).

The prepared cDNA libraries were sequenced on a MiSeq (Illumina). Reads from each sample were identified on the basis of their associated barcode and aligned to the appropriate RefSeq reference genome using BWA (Li and Durbin, 2009). Paired-end alignments were used to extract entire transcript sequences using Galaxy (https://usegalaxy.org), and these sequences were analyzed using Geneious 8.1.8.

### *E. coli* Essential Gene Screen Experiment

The intersection of two *E. coli* DH10B strain essential gene studies (Gerdes et al., 2003; Baba et al., 2006) was taken, and further pared down to 45 genes by only selecting genes exclusive to their respective operons (Table S2). Over these 45 genes 54,600 spacers were designed to tile at single resolution across the coding region, as well as to extend 60nt into the 5’ UTR and 3’UTR. In addition, 1100 non-targeting, pseudorandomly generated spacers with no precise match to the *E. coli* DH10B strain genome were added to the library as a non-targeting negative control. The library of spacers (Table S3) was cloned into a *B. zoohelcum* or *P. buccae* direct repeat-spacer-direct repeat backbone containing a chloramphenicol resistance gene using Golden Gate Assembly (NEB) with 100 cycles, and then transformed over five 22.7cm x 22.7cm chloramphenicol LB Agar plates. Libraries of transformants were scraped from plates and DNA was extracted using the Macherey-Nagel Nucleobond Xtra Midiprep Kit (Macherey-Nagel). 50ng of library plasmid and equimolar gene plasmid containing an ampicillin resistance gene *(bzcas13b, bzcas13b & bzcsx27, pbcas13b, pbcas13b & pbcsx28,* empty vector pBR322 (Table S1)) were transformed into MegaX DH10B^TM^ T1R Electrocomp^TM^ Cells (ThermoFisher) according to manufacturer’s protocol, with four separate 22.7cm x 22.7cm carbenicillin-chloramphenicol LB Agar plates per bioreplicate, and three bioreplicates per condition (twelve transformations total per condition). Eleven hours post-transformation, libraries of transformants were scraped from plates and DNA extracted using the Macherey-Nagel Nucleobond Xtra Maxiprep Kit (Macherey-Nagel).

### *E. coli* Essential Gene Screen Analysis

Prepared DNA libraries were sequenced on a NextSeq (Illumina), with reads mapped to the input library of spacers. Spacer depletions were calculated as the read abundance of a spacer in the empty vector condition divided by read abundance in each gene plasmid condition. Mean depletions over three bioreplicates were calculated. We imposed a two-step quality-control filter on the data: a maximum coefficient of variation of 0.2 for depletion over three bioreplicates, and a minimum spacer read abundance of 1/3*N* in each bioreplicate, where N = 55,700. Weblogos of the strongly depleted (top 1% depleted) spacers were generated (Crooks et al., 2004), and from each identified PFS, heatmaps of the ratio of safely depleted (>5σ above mean depletion of nontargeting spacers) spacers to all spacers in the screen were generated. For spatial analysis via empirical cumulative distribution functions, safely depleted spacers were aggregated across the first or last 250nt of genes.

For secondary structure analysis, we utilized the RNA accessibility model from Vienna RNAplfold (Bernhart et al., 2006). RNAplfold calculates through a moving average of RNA folds the probability that a region *u* of RNA is unpaired given its *cis* sequence context in a four-parameter model, where *W* is the moving average window length in nucleotides, *L* is the maximum permissible pairing distance between nucleotides in the window, and *u_start_* and *u_end_* are the start and end of the region *u*, respectively. To apply this model to our data, we separated spacers from our *E. coli* essential gene screen into training/testing cohorts of five or more, each represented by a unique permissible PFS and gene and containing at least one spacer in the top 2% of depleted spacers from the screen (to enhance predictive signal). We then randomly divided these cohorts into a training set (~80%) and a testing set (~20%). For optimizing a secondary structure-mediated model of efficient spacer design we selected as objective functions *top 1* or *top 3 accuracy,* the percent of cohorts for which the top spacer is accurately predicted or falls in the top 3 depleted spacers in a cohort, respectively. We optimized the two objective functions on the training data set, first by fixing *W* and *L* while varying *u_start_* and *u_end_*, then by fixing *u_start_* and *u_end_* and varying *W* and *L* (Figure 4B). In the case of *bzcas13b* with *bzcsx27*, as well as that of *bzcas13b* alone, the optimized parameters were found to be approximately *W* = 240, *L* = 180, *u_start_* = 16, and *u_end_* = 30. We gauged the performance of this RNAplfold model relative to 10^6^ Monte Carlo simulations performed on the testing data set and found empirical *P-* values of less than 1e-2 for *top 1 accuracy*, and less than 1e-5 for *top 3 accuracy*. Similar predictive power applied to *pbcas13b* with *pbcsx28*, as well as to *pbcas13b* alone.

### Kanamycin Validation Screen Experiment

A total of 160 kanamycin-targeting spacers was selected, 42 of which contain both PFS rules, 47 of which contain one rule, and 71 of which contain no rules, to which 162 non-targeting control spacers were added (Table S4). The library of spacers was cloned into either a *bzcas13b* and *B. zoohelcum* direct repeat-spacer-direct repeat backbone or simply a *B. zoohelcum* direct repeat-spacer-direct repeat backbone containing a chloramphenicol resistance gene using Golden Gate Assembly (NEB) with 100 cycles, and then transformed over one 22.7cm × 22.7cm carbenicillin LB Agar plate. The two cloned library plasmids were then re-transformed with over a 22.7cm × 22.7cm chloramphenicol LB Agar plate or a 22.7cm × 22.7cm kanamycin-chloramphenicol LB Agar plate. Libraries of transformants were scraped from plates and DNA extracted using the Qiagen Plasmid Plus Maxi Kit (Qiagen). 100 ng of library DNA and 100 ng of pMAX-GFP (Lonza), containing a kanamycin resistance gene were added to 50 uL of chemically competent 10-beta cells (NEB) and transformed according to the manufacturer’s protocol.

### Kanamycin Validation Screen Analysis

Prepared DNA libraries were sequenced on a NextSeq (Illumina), with reads mapped to the input library of spacers. For normalizing the abundance of spacers of two separate clonings, the corrected experimental read abundance of a given spacer was calculated as the read abundance of that spacer in the *bzcas13b* plasmid (kanamycin-chloramphenicol transformation) multiplied by the ratio of the read abundance ratio of that spacer in the non-*bzcas13b* plasmid (chloramphenicol-only transformation) to the read abundance ratio of that spacer in the *bzcas13b* plasmid (chloramphenicol-only transformation).

### MS2 Phage Drop Plaque Assay

Individual spacers for MS2 interference were ordered as complementary oligonucleotides containing overhangs allowing for directional cloning in between two direct repeat sequences in vectors containing *cas13b* (Tables S1, S5). 10 uM of each complementary oligo were annealed in 10X PNK Buffer (NEB), supplemented with 10mM ATP and 5 units of T4PNK (NEB). Oligos were incubated at 37C for 30 min., followed by heating to 95C for 5 min. and then annealed by cooling to 4C. Annealed oligos were then diluted 1:100 and incubated with 25 ng of Eco31I digested *cas13b* vector in the presence of Rapid Ligation Buffer and T7 DNA ligase (Enzymatics). Individual plasmids were prepared using the QIAprep Spin Miniprep Kit (Qiagen), sequence confirmed and then transformed into C3000 (ATCC 15597) cells made competent using the Mix & Go *E. coli* Transformation Kit (Zymo). In the case of experiments using *csx27* or *csx28,* C3000 cells harboring these plasmids were made competent and then transformed with *cas13b* direct repeat-spacer-direct repeat plasmids. Following transformation, individual clones were picked and grown overnight in LB containing the appropriate antibiotics. The following morning, cultures were diluted 1:100 and grown to an OD_600_ of 2.0, then mixed with 4mL of antibiotic containing Top Agar (10 g/L tryptone, 5 g/L yeast extract, 10 g/L sodium chloride, 5 g/L agar) and poured on to LB-antibiotic base plates. 10 fold serial-dilutions of MS2 phage were made in LB and then spotted onto hardened top agar with a multi-channel pipette. For assessing interference levels in Figures 5E and 7D, samples were blinded using a key and the lowest dilution of phage at which plaque formation occurred was compared to a pACYC condition by eye, where the lowest dilution of MS2 that formed plaques on pACYC was set to 1. The lowest dilution of phage used for Figure 5E was 1.05^*^10^8^ pfu.

### DNA Interference Assay

A 34-nt target sequence consisting of a 30-nt protospacer and a permissive PFS (5’-G, 3’-AAA) was cloned into pUC19 in two locations (Tables S1, S5). For the transcribed target, the target sequence was cloned into the coding strand of the *bla* gene, in frame immediately after the start codon, with the G of the start codon serving as the 5’ PFS. For the non-transcribed target the identical target sequence (protospacer and PFS) were cloned into the AatII site of pUC19, so that the protospacer appears on the non-transcribed strand with respect to the pBla and pLac promoters. To determine interference, 25 ng of the ampicillin resistant target plasmid and 25 ng of the chloramphenicol resistant *bzcas13b* or empty vector (pACYC) were added to 5 uL of NovaBlue GigaSingle cells (Novagen). The cells were incubated for 30 minutes on ice, heatshocked for 30 seconds at 42°C and incubated on ice for 2 minutes. Then, 95 uL of SOC was added to cells and they were incubated with shaking at 37°C for 90 minutes, before plating the entire outgrowth (100 uL) on plates containing both chloramphenicol and ampicillin.

### BzCas13b Protein Purification

The mammalian codon-optimized gene for Cas13b *(B. zoohelcum)* was synthesized (GenScript) and inserted into a bacterial expression vector (6× His/Twin Strep SUMO, a pET based vector received as a gift from Ilya Finkelstein) after cleaving the plasmid with the BamHI and NotI restriction enzymes and cloning in the gene using Gibson Assembly® Master Mix (New England Biolabs). The BzCas13b expression construct (Table S1) was transformed into One Shot® BL21(DE3)pLysE (Invitrogen) cells. 25 mL of 6hr growing culture were inoculated into 2 liters of Terrific Broth 4 growth media (12 g/L tryptone, 24 g/L yeast extract, 9.4 g/L K2HPO, 2.2 g/L KH2PO4, Sigma). Cells were then grown at 37°C to a cell density of 0.6 OD600, and then SUMO-BzCas13b expression was induced by supplementing with IPTG to a final concentration of 500 μM. Induced culture was grown for 16-18 hours before harvesting cell paste, which was stored at −80ºC until subsequent purification. Frozen cell paste was crushed and resuspended via stirring at 4°C in 500 mL of Lysis Buffer (50mM NaH_2_PO_4_ pH 7.8, 400mM NaCl) supplemented with protease inhibitors (cOmplete, EDTA-free, Roche Diagnostics Corporation) and 1250U of benzonase (Invitrogen). The resuspended cell paste was lysed by a LM20 microfluidizer at 18,000 psi (Microfluidics). Lysate was cleared by centrifugation at 10,000g for 1 hour. Filtered lysate was incubated with StrepTactin Sepharose High Performance (GE Healthcare Life Sciences) at 4°C for 1 hour with gentle agitation, and then applied to an Econo-column chromatography column (Bio-Rad Laboratories). Resin was washed with Lysis Buffer for 10 column volumes. One column volume of fresh Lysis Buffer was added to the column and mixed with 10 units of SUMO protease (Invitrogen) and incubated overnight. The eluate was removed from the column, SUMO cleavage was confirmed by SDS-PAGE and BlueFast protein staining (Eton Bioscience), and the sample was concentrated via Centrifugal Filter Unit to 2 mL. Concentrated sample was loaded onto a HiTrap Heparin HP column (GE Healthcare Life Sciences) via FPLC (AKTA Pure, GE Healthcare Life Sciences) and eluted over a gradient with an elution buffer with salt concentration of 1.2 M. The resulting fractions were tested for presence of BzCas13b protein by SDS-PAGE; fractions containing BzCas13b were pooled, and concentrated via Centrifugal Filter Unit to 1 mL. Concentrated sample was loaded a gel filtration column (HiLoad 16/600 Superdex 200, GE Healthcare Life Sciences) via FPLC (AKTA Pure, GE Healthcare Life Sciences) with buffer 500 mM NaCl, 50 mM Tris-HCl pH 7.5, 1 mM DTT.

### BzCas13b HEPN Mutant Protein Purification

Alanine mutants (Table S1) at each of the HEPN catalytic residues were generated using the Q5^®^ site-directed mutagenesis kit (New England Biolabs) and transformed into One ShotⓇ BL21(DE3)pLysE cells (Invitrogen). For each mutant, 1 L of Terrific Broth was used to generate cell paste and all other reagents were scaled down accordingly. Protein purification was performed using the same protocol as wild-type Cas13b.

### PbCas13b Protein Purification

PbCas13b *(Prevotella buccae)* was cloned into the same pET based vector and purified using a similar protocol as BzCas13b with the following differences: cells were grown at 21°C for 18 hours. Frozen cell paste was resuspended into 500 mM NaCl, 50 mM HEPES 7.5 and 2 mM DTT prior to breaking cells in the microfluidizer. The Superdex 200 column was run in 500 mM NaCl, 10 mM HEPES 7.0, and 2 mM DTT.

### Nucleic Acid Preparation

For in vitro synthesis of RNA, a T7 DNA fragment must be generated. To create T7 DNA fragments for crRNAs, top and bottom strand DNA oligos were synthesized by IDT. The top DNA oligo consisted of the T7 promoter, followed by the bases GGG to promote transcription, the 30-nt target and then direct repeat. Oligos were annealed together using annealing buffer (30 mM HEPES pH 7.4, 100 mM potassium acetate, and 2 mM magnesium acetate). Annealing was performed by incubating the mixture for 1 minute at 95°C followed by a −1°C/minute ramp down to 23°C. To create ssRNA targets, short targets (Trunc2, 3, 4) were synthesized as top and bottom strand oligos containing the T7 promoter. For long ssRNA targets (E1, E2, S and L CRISPR Arrays), DNA primers (Table S8) with a T7 handle on the forward primer were ordered and the DNA fragment was amplified using PCR. T7 DNA constructs for RNA generation without body labeling were incubated with T7 polymerase overnight (10-14 hours) at 30°C using the HiScribe T7 Quick High Yield RNA Synthesis kit (New England Biolabs). Body-labeled constructs were incubated with Cyanine 5-UTP (Perkin Elmer) and incubated with T7 polymerase overnight at 30°C using the HiScribe T7 High Yield RNA Synthesis kit (New England Biolabs). For a complete list of crRNAs used in this study see Table S6. For a complete list of targets used in this study see table S7. 5’ end labeling was accomplished using the 5’ oligonucleotide kit (VectorLabs) and with a maleimide-IR800 probe (LI-COR Biosciences). 3’ end labeling was performed using a 3’ oligonucleotide labeling kit (Roche) and Cyanine 5-ddUTP (Perkin Elmer). RNAs were purified using RNA Clean and Concentrator columns^TM^-5(Zymo Research). Body-labeled dsRNA substrates were prepared by T7 DNA fragments for the bottom and top RNA strand. After synthesis, 1.3-fold excess of non-labeled bottom strand ssRNA was added and re-annealed to ensure the top strand would be annealed to a bottom strand by incubating the mixture for 1 minute at 95°C followed by a −1°C/minute ramp down to 23°C.

### Nuclease Assay

Nuclease assays were performed with equimolar amounts of end-labeled or body-labeled ssRNA target, purified protein, and crRNA, for targeted ssRNA cleavage. For CRISPR array cleavage, protein was supplied in a four times molar excess of the CRISPR array. Reactions were incubated in nuclease assay buffer (10 mM TrisHCl pH 7.5, 50 mM NaCl, 0.5 mM MgCl_2_, 0.1% BSA). Reactions were allowed to proceed at 37°C for times specified in the figure legends. After incubation, samples were then quenched with 0.8U of Proteinase K (New England Biolabs) for 15 minutes at 25°C. The reactions were mixed with equal parts of RNA loading dye (New England Biolabs) and denatured at 95°C for 5 minutes and then cooled on ice for 2 minutes. Samples were analyzed by denaturing gel electrophoresis on 10% PAGE TBE-Urea (Invitrogen) run at 45°C. Gels were imaged using an Odyssey scanner (LI-COR Biosciences).

### EMSA Assay

For the Electrophoretic Mobility Shift Assay (EMSA), binding experiments were performed with a series of half-log complex dilutions (crRNA and BzCas13b) from .594 to 594 nM. Binding assays were performed in nuclease assay buffer supplemented with 10 mM EDTA to prevent cutting, 5% glycerol, and 5μg/mL heparin in order to avoid non-specific interactions of the complex with target RNA. Protein was supplied at two times the molar amount of crRNA. Protein and crRNA were preincubated at 37°C for 15 minutes, after which the 5’-labeled target was added. Reactions were then incubated at 37°C for 10 minutes and then resolved on 6% PAGE TBE gels (Invitrogen) at 4°C (using 0.5X TBE buffer). Gels were imaged using an Odyssey scanner (LI-COR Biosciences). Gel shift of the targets was quantified using ImageJ and plotted in Prism7 (GraphPad). Line regression was performed in Prism 7 using nonlinear fit and specific binding with Hill slope.

## Supplementary Figures and Tables

**Figure S1 | Phylogenetic tree of Cas13b bifurcates into two variants of subtype VI-B CRISPR loci.** A phylogenetic tree (alignment generated by BLOSUM62) of non-redundant Cas13b effectors, with the full type VI-B locus depicted in every instance. Accession numbers for genome, Cas13b (blue), and Csx27 (brown)/Csx28 (gold) are included, as well as number of nearby spacers detected by PILER-CR, the presence of other CRISPR-Cas elements in the genome, and the size of Cas13b.

**Figure S2 | HEPN domains in Cas13b and Csx28 from multiple sequence alignments. (A)** Two HEPN sequences identified via multiple sequence alignment (BLOSUM62) of putative non-redundant Cas13b proteins. **(B)** Divergent HEPN sequence identified via multiple sequence alignment (BLOSUM62) of putative non-redundant Csx28 proteins.

**Figure S3 | Predicted transmembrane domains of Csx27 and Csx28 not validated experimentally. (A)** Transmembrane domain prediction in Csx27 of *B. zoohelcum* and Csx28 of *P. buccae* using TMHMM v2. **(B)** N- and C-terminally fused RFP imaging of Csx27 of *B. zoohelcum* and Csx28 of *P. buccae.*

**Figure S4 | Predicted secondary structure of type VI-B direct repeats is well-conserved. (A)** Predicted secondary structure folds of structurally unique CRISPR class 2 type VI-B1 direct repeats (Vienna RNAfold). **(B)** Predicted secondary structure folds of structurally unique CRISPR Class 2 type VI-B2 direct repeats.

**Figure S5 | Type VI-B direct repeats are well-conserved; predicted protospacer flanking sequences are inconclusive. (A)** Weblogo of all unique VI-B direct repeat sequences of length 36nt, taken as the same transcriptional orientation as Cas13b. **(B)** Weblogo of all unique VI-B protospacer flanking sequences from CRISPRTarget mapping of protospacers to phage databases.

**Figure S6 | Protein gels of purified WT BzCas13b and three mutant BzCas13b proteins.** Denaturing protein gels of *B. zoohelcum* wildtype, R116A/H121A mutant, R1177A/H1182A mutant, and R116A/H121A/R1177A/H1182A mutant Cas13b.

**Figure S7 | Second bioreplicate of kanamycin validation screen for Cas13b from *B. zoohelcum* agrees with first. (A)** Spacers targeting kanamycin to validate PFS targeting rules of 5’ PFS (D) and 3’ PFS (NAN or NNA). **(B)** Spacer abundances versus control for individual *B. zoohelcum* spacers, with abundances colored by type of spacer.

**Figure S8 | Cas13b cleaves single-stranded RNA. (A)** Denaturing gels demonstrating no cleavage of dsRNA, ssDNA, or dsDNA by BzCas13b with either the short DR or long DR. Reactions were incubated for 10 minutes, the same amount of time which results in robust ssRNA cleavage for this target and crRNA pair. The ssDNA and top strand of the dsDNA target is 5′ labeled with IRDye 800. The dsRNA target is body labeled. **(B)** ssRNA cleavage requires BzCas13b and a targeting crRNA and this cleavage activity is abolished by addition of EDTA. **(C)** Denaturing gel showing PbCas13b cleavage activity of an ssRNA targeted substrate. The ssRNA is 5′ labeled with IRDye 800 and incubated for 30 minutes.

**Figure S9 | Bioreplicates of MS2 phage plaque drop assay.** Plaque drop assay with bioreplicates for *B. zoohelcum* VI-B1 locus and *cas13b,* for *P. buccae* VI-B2 locus and *cas13b,* and for *P. buccae cas13b* with pUC19, *B. zoohelcum csx27,* and *P. buccae csx28.*

**Figure S10 | HEPN mutant protein and RNA binding. (A)** EMSA gels that were used to quantify the K_D_ of the WT and mutant BzCas13b proteins, using an on-target crRNA complementary to the targeted ssRNA. **(B)** EMSA gel of WT BzCas13b with an off-target crRNA. The off-target crRNA is non-complementary to the targeted ssRNA.

**Figure S11 | RNA-Sequencing of *P. buccae* VI-B2 CRISPR locus.** RNA-Sequencing of heterologously expressed VI-B2 locus from *P. buccae* ATCC 33574 in *E. coli.*

**Figure S12 | *E. coli* essential gene screen of *P. buccae* VI-B2 CRISPR locus. (A)** Manhattan plots of spacer depletions mapped over 45 genes and aggregated across normalized gene distance for full *P. buccae* VI-B2 locus (left) and *cas13b* (right), with non-targeting spacers in gray, safely depleted (>5σ above mean depletion of non-targeting spacers) spacers above blue line, and strongly depleted (top 1% depleted) spacers above red line. For the full locus, 36,141 targeting spacers and 859 non-targeting spacers passed QC filter. Of the targeting, 370 are strongly depleted and 8065 are safely depleted. For *cas13b* alone, 41,126 targeting spacers and 824 non-targeting spacers passed QC filter. Of the targeting, 419 are strongly depleted and 3295 are safely depleted. **(B)** Sequence weblogos of strongly depleted *P. buccae* spacers, revealing double-sided PFS (protospacer flanking sequence).

**Figure S13 | Computational analysis of secondary structure and spatial rules for *P. buccae cas13b* RNA targeting. (A)** Optimization of *top 1 accuracy* (computationally predicted spacer is top depleted) and *top 3 accuracy* (computationally predicted spacer falls in top 3 depleted) on randomly selected *P. buccae* training dataset using RNAplfold, first with *u start* and *u end,* and then with *W* and *L.* **(B)** Performance of optimized RNAplfold model on randomly selected *P. buccae* testing dataset (41 cohorts for full *P. buccae* VI-B2 locus, 40 cohorts for *pbcas13b*) against 10^6^ Monte Carlo simulations: empirical P-values from left to right of 3.3e-2, 2.7e-3, 3.9e-3, 1.5e-5. **(C)** Empirical cumulative distribution function of safely depleted *P. buccae* spacers over all genes from 5’UTR into gene and from 3’ UTR into gene. Yellow line separates UTR and gene, red line is theoretical cumulative distribution function of uniformly distributed spacers, and blue line is empirical cumulative distribution of safely depleted *P. buccae* spacers.

**Figure S14 | A differential regulatory model for VI-B CRISPR systems.** VI-B systems encode the RNA-guide RNase Cas13b, repressed by Csx27 and enhanced by Csx28, whose RNA interference is governed by a sequence-and-structure RNA targeting model.

**Table S1 | All Cas13b plasmids used in this study. Related to Figures 2, 3, 5, S5, and S9.**

**Table S2 | *E. coli* essential genes represented in *E. coli* essential gene screen library of spacers.**

**Table S3 | Spacers from *E. coli* essential gene screen.**

**Table S4 | Spacers from kanamycin validation screen.**

**Table S5 | Spacers targeting MS2 and pBLA plasmids.**

**Table S6 | All crRNAs used in biochemical experiments.**

**Table S7 | All nucleic acid targets used in biochemical experiments.**

**Table S8 | T7 primers used to generate long single-stranded RNA targets.**

## References

1. Abil Z, Zhao H. Engineering reprogrammable RNA-binding proteins for study and manipulation of the transcriptome. Molecular bioSystems. 2015; 11(10):2658–65. doi: 10.1039/c5mb00289c. PubMed PMID: 26166256.

2. Abudayyeh OO, Gootenberg JS, Konermann S, Joung J, Slaymaker IM, Cox DB, et al. C2c2 is a single-component programmable RNA-guided RNA-targeting CRISPR effector. Science. 2016;353(6299):aaf5573. doi: 10.1126/science.aaf5573. PubMed PMID: 27256883.

3. Anantharaman V, Makarova KS, Burroughs AM, Koonin EV, Aravind L. Comprehensive analysis of the HEPN superfamily: identification of novel roles in intra-genomic conflicts, defense, pathogenesis and RNA processing. Biology direct. 2013;8:15. doi: 10.1186/17456150-8-15. PubMed PMID: 23768067; PubMed Central PMCID: PMC3710099.

4. Baba T, Ara T, Hasegawa M, Takai Y, Okumura Y, Baba M, et al. Construction of Escherichia coli K-12 in-frame, single-gene knockout mutants: the Keio collection. Molecular systems biology. 2006;2:2006–2008. doi: 10.1038/msb4100050. PubMed PMID: 16738554; PubMed Central PMCID: PMC1681482.

5. Bernhart SH, Hofacker IL, Stadler PF. Local RNA base pairing probabilities in large sequences. Bioinformatics. 2006;22(5):614–5. doi: 10.1093/bioinformatics/btk014. PubMed PMID: 16368769.

6. Biswas A, Gagnon JN, Brouns SJ, Fineran PC, Brown CM. CRISPRTarget: bioinformatic prediction and analysis of crRNA targets. RNA biology. 2013;10(5):817–27. doi: 10.4161/rna.24046. PubMed PMID: 23492433; PubMed Central PMCID: PMC3737339.

7. Camacho C, Coulouris G, Avagyan V, Ma N, Papadopoulos J, Bealer K, et al. BLAST+: architecture and applications. BMC bioinformatics. 2009;10:421. doi: 10.1186/1471-210510-421. PubMed PMID: 20003500; PubMed Central PMCID: PMC2803857.

8. Cong L, Ran FA, Cox D, Lin S, Barretto R, Habib N, et al. Multiplex genome engineering using CRISPR/Cas systems. Science. 2013;339(6121):819–23. doi: 10.1126/science.1231143. PubMed PMID: 23287718; PubMed Central PMCID: PMC3795411.

9. Crooks GE, Hon G, Chandonia JM, Brenner SE. WebLogo: a sequence logo generator. Genome research. 2004;14(6):1188–90. doi: 10.1101/gr.849004. PubMed PMID: 15173120; PubMed Central PMCID: PMC419797.

10. East-Seletsky A, O’Connell MR, Knight SC, Burstein D, Cate JH, Tjian R, et al. Two distinct RNase activities of CRISPR-C2c2 enable guide-RNA processing and RNA detection. Nature. 2016. doi: 10.1038/nature19802. PubMed PMID: 27669025.

11. Edgar RC. PILER-CR: fast and accurate identification of CRISPR repeats. BMC bioinformatics. 2007;8:18. doi: 10.1186/1471-2105-8-18. PubMed PMID: 17239253; PubMed Central PMCID: PMC1790904.

12. Filipovska A, Rackham O. Designer RNA-binding proteins: New tools for manipulating the transcriptome. RNA biology. 2011;8(6):978–83. doi: 10.4161/rna.8.6.17907. PubMed PMID: 21941129.

13. Gerdes SY, Scholle MD, Campbell JW, Balazsi G, Ravasz E, Daugherty MD, et al. Experimental determination and system level analysis of essential genes in Escherichia coli MG1655. Journal of bacteriology. 2003;185(19):5673–84. PubMed PMID: 13129938; PubMed Central PMCID: PMC193955.

14. Hale CR, Cocozaki A, Li H, Terns RM, Terns MP. Target RNA capture and cleavage by the Cmr type III-B CRISPR-Cas effector complex. Genes & development. 2014;28(21):2432–43. doi: 10.1101/gad.250712.114. PubMed PMID: 25367038; PubMed Central PMCID: PMC4215187.

15. Hale CR, Majumdar S, Elmore J, Pfister N, Compton M, Olson S, et al. Essential features and rational design of CRISPR RNAs that function with the Cas RAMP module complex to cleave RNAs. Molecular cell. 2012;45(3):292–302. doi: 10.1016/j.molcel.2011.10.023. PubMed PMID: 22227116; PubMed Central PMCID: PMC3278580.

16. Hale CR, Zhao P, Olson S, Duff MO, Graveley BR, Wells L, et al. RNA-guided RNA cleavage by a CRISPR RNA-Cas protein complex. Cell. 2009;139(5):945–56. doi: 10.1016/j.cell.2009.07.040. PubMed PMID: 19945378; PubMed Central PMCID: PMC2951265.

17. Hayes F, Van Melderen L. Toxins-antitoxins: diversity, evolution and function. Critical reviews in biochemistry and molecular biology. 2011;46(5):386–408. doi: 10.3109/10409238.2011.600437. PubMed PMID: 21819231.

18. Heidrich N, Dugar G, Vogel J, Sharma CM. Investigating CRISPR RNA Biogenesis and Function Using RNA-seq. Methods in molecular biology. 2015;1311:1–21. doi: 10.1007/978-1-4939-2687-9_1. PubMed PMID: 25981463.

19. Henikoff S, Henikoff JG. Amino acid substitution matrices from protein blocks. Proceedings of the National Academy of Sciences of the United States of America. 1992;89(22):10915–9. PubMed PMID: 1438297; PubMed Central PMCID: PMC50453.

20. Jiang W, Samai P, Marraffini LA. Degradation of Phage Transcripts by CRISPR-Associated RNases Enables Type III CRISPR-Cas Immunity. Cell. 2016;164(4):710–21. doi: 10.1016/j.cell.2015.12.053. PubMed PMID: 26853474; PubMed Central PMCID: PMC4752873.

21. Kim YK, Kim YG, Oh BH. Crystal structure and nucleic acid-binding activity of the CRISPR-associated protein Csx1 of Pyrococcus furiosus. Proteins. 2013;81(2):261–70. doi: 10.1002/prot.24183. PubMed PMID: 22987782.

22. Li H, Durbin R. Fast and accurate short read alignment with Burrows-Wheeler transform. Bioinformatics. 2009;25(14):1754–60. doi: 10.1093/bioinformatics/btp324. PubMed PMID: 19451168; PubMed Central PMCID: PMC2705234.

23. Lorenz R, Bernhart SH, Honer Zu Siederdissen C, Tafer H, Flamm C, Stadler PF, et al. ViennaRNA Package 2.0. Algorithms for molecular biology: AMB. 2011;6:26. doi: 10.1186/1748-7188-6-26. PubMed PMID: 22115189; PubMed Central PMCID: PMC3319429.

24. Mackay JP, Font J, Segal DJ. The prospects for designer single-stranded RNA-binding proteins. Nature structural & molecular biology. 2011;18(3):256–61. doi: 10.1038/nsmb.2005. PubMed PMID: 21358629.

25. Makarova KS, Anantharaman V, Aravind L, Koonin EV. Live virus-free or die: coupling of antivirus immunity and programmed suicide or dormancy in prokaryotes. Biology direct. 2012;7:40. doi: 10.1186/1745-6150-7-40. PubMed PMID: 23151069; PubMed Central PMCID: PMC3506569.

26. Makarova KS, Wolf YI, Alkhnbashi OS, Costa F, Shah SA, Saunders SJ, et al. An updated evolutionary classification of CRISPR-Cas systems. Nature reviews Microbiology. 2015;13(11):722–36. doi: 10.1038/nrmicro3569. PubMed PMID: 26411297.

27. Makarova KS, Wolf YI, Koonin EV. Comprehensive comparative-genomic analysis of type 2 toxin-antitoxin systems and related mobile stress response systems in prokaryotes. Biology direct. 2009;4:19. doi: 10.1186/1745-6150-4-19. PubMed PMID: 19493340; PubMed Central PMCID: PMC2701414.

28. Mali P, Yang L, Esvelt KM, Aach J, Guell M, DiCarlo JE, et al. RNA-guided human genome engineering via Cas9. Science. 2013;339(6121):823–6. doi: 10.1126/science.1232033. PubMed PMID: 23287722; PubMed Central PMCID: PMC3712628.

29. Marraffini LA. CRISPR-Cas immunity in prokaryotes. Nature. 2015;526(7571):55–61. doi: 10.1038/nature15386. PubMed PMID: 26432244.

30. Mohanraju P, Makarova KS, Zetsche B, Zhang F, Koonin EV, van der Oost J. Diverse evolutionary roots and mechanistic variations of the CRISPR-Cas systems. Science. 2016;353(6299):aad5147. doi: 10.1126/science.aad5147. PubMed PMID: 27493190.

31. Moller S, Croning MD, Apweiler R. Evaluation of methods for the prediction of membrane spanning regions. Bioinformatics. 2001;17(7):646–53. PubMed PMID: 11448883.

32. Ran FA, Cong L, Yan WX, Scott DA, Gootenberg JS, Kriz AJ, et al. In vivo genome editing using Staphylococcus aureus Cas9. Nature. 2015;520(7546):186–91. doi: 10.1038/nature14299. PubMed PMID: 25830891; PubMed Central PMCID: PMC4393360.

33. Remmert M, Biegert A, Hauser A, Soding J. HHblits: lightning-fast iterative protein sequence searching by HMM-HMM alignment. Nature methods. 2012;9(2):173–5. doi: 10.1038/nmeth.1818. PubMed PMID: 22198341.

34. Sheppard NF, Glover CV, 3rd, Terns RM, Terns MP. The CRISPR-associated Csx1 protein of Pyrococcus furiosus is an adenosine-specific endoribonuclease. Rna. 2016;22(2):216–24. doi: 10.1261/rna.039842.113. PubMed PMID: 26647461; PubMed Central PMCID: PMC4712672.

35. Shmakov S, Abudayyeh OO, Makarova KS, Wolf YI, Gootenberg JS, Semenova E, et al. Discovery and Functional Characterization of Diverse Class 2 CRISPR-Cas Systems. Molecular cell. 2015;60(3):385–97. doi: 10.1016/j.molcel.2015.10.008. PubMed PMID: 26593719; PubMed Central PMCID: PMC4660269.

36. Shmakov S, Smargon A, Scott D, Cox D, Pyzocha N, Yan W, et al. Diversity and evolution of Class 2 CRISPR-Cas systems. Nature reviews microbiology. *In press.*

37. Staals RH, Agari Y, Maki-Yonekura S, Zhu Y, Taylor DW, van Duijn E, et al. Structure and activity of the RNA-targeting Type III-B CRISPR-Cas complex of Thermus thermophilus. Molecular cell. 2013;52(1): 135–45. doi: 10.1016/j.molcel.2013.09.013. PubMed PMID: 24119403; PubMed Central PMCID: PMC4006948.

38. Staals RH, Zhu Y, Taylor DW, Kornfeld JE, Sharma K, Barendregt A, et al. RNA targeting by the type III-A CRISPR-Cas Csm complex of Thermus thermophilus. Molecular cell. 2014;56(4):518–30. doi: 10.1016/j.molcel.2014.10.005. PubMed PMID: 25457165; PubMed Central PMCID: PMC4342149.

39. Tafer H, Ameres SL, Obernosterer G, Gebeshuber CA, Schroeder R, Martinez J, et al. The impact of target site accessibility on the design of effective siRNAs. Nature biotechnology. 2008;26(5):578–83. doi: 10.1038/nbt1404. PubMed PMID: 18438400.

40. Tamulaitis G, Kazlauskiene M, Manakova E, Venclovas C, Nwokeoji AO, Dickman MJ, et al. Programmable RNA shredding by the type III-A CRISPR-Cas system of Streptococcus thermophilus. Molecular cell. 2014;56(4):506–17. doi: 10.1016/j.molcel.2014.09.027. PubMed PMID: 25458845.

41. Wright AV, Nunez JK, Doudna JA. Biology and Applications of CRISPR Systems: Harnessing Nature’s Toolbox for Genome Engineering. Cell. 2016;164(1-2):29–44. doi: 10.1016/j.cell.2015.12.035. PubMed PMID: 26771484.

42. Yates A, Akanni W, Amode MR, Barrell D, Billis K, Carvalho-Silva D, et al. Ensembl 2016. Nucleic acids research. 2016;44(D1):D710–6. doi: 10.1093/nar/gkv1157. PubMed PMID: 26687719; PubMed Central PMCID: PMC4702834.

43. Zhang J, Rouillon C, Kerou M, Reeks J, Brugger K, Graham S, et al. Structure and mechanism of the CMR complex for CRISPR-mediated antiviral immunity. Molecular cell. 2012;45(3):303–13. doi: 10.1016/j.molcel.2011.12.013. PubMed PMID: 22227115; PubMed Central PMCID: PMC3381847.

